# Persistent hindrances to data re-use in single-cell genomics

**DOI:** 10.1101/2025.10.02.680150

**Authors:** Sanja Rogic, Xinrui Xiang Yu, Brianna Xu, Alexandra Millett, Salva Sherif, Guillaume Poirier-Morency, Rachel Schwartz, Paul Pavlidis

## Abstract

We report on our experience attempting to re-use published and publicly available single-cell (or single-nucleus) RNA-sequencing studies (scRNA-seq) from the Gene Expression Omnibus (GEO). We screened GEO for human, mouse and rat scRNA-seq studies as potential candidates for inclusion in the Gemma database of re-annotated and re-analyzed transcriptome studies. Using semi-automated and manual curation, we assessed whether GEO datasets included cell-level expression count matrices and cell-type annotations. We found that there are steep challenges to data reuse. Only ∼40% of studies provided readily usable processed count data that could be reliably mapped to GEO metadata, and fewer than 10% included author-provided cell-type annotations. While raw sequencing data were available for the majority of studies, only a small proportion could be re-analyzed automatically without reliance on heuristics. Our findings show that existing practices for single-cell RNA-sequencing data distribution and sharing are insufficient for effective reuse, and highlight the urgent need for repositories to strengthen and enforce submission requirements, particularly for processed data and cell-type annotations.

## Introduction

Data re-use has become a bedrock of genomics research, enabled by open sharing of data through resources such as the Gene Expression Omnibus (GEO) (Clough et al., 2023). However, in practice there are hurdles to re-use, some of which we have documented in our previous work, and which we attempt to address through re-curation in our Gemma resource (https://gemma.msl.ubc.ca; (Lim et al., 2021)). General challenges include the lack of standardization (or even absence) of experimental design information, lack of detailed platform information (for microarrays), as well as various data quality issues.

Single-cell data opens up a new set of challenges to re-use. While access to count-level expression data is equally important for “traditional” (bulk) RNA-seq, for scRNAseq there are more variables to consider if attempting to recreate them from raw sequence data. The predominant platform for single-cell transcriptomics is 10x Chromium, accounting for approximately 80% of single-cell datasets in GEO. 10x provides proprietary (but free of cost) software (Cell Ranger, https://github.com/10XGenomics/cellranger) that handles the initial stages of analysis of the data, in particular, de-multiplexing of cells and quantification of gene expression. Having such data provided by the authors of each study would greatly simplify attempts to reproduce or extend the study, because most re-users would not want to repeat these steps without a good reason, especially given the potential for difficulty in reproducing the exact methods of the study originators.

Another essential component of a single-cell dataset is the annotation of each cell in the study. Essentially every publication that uses single-cell data assigns cells to “types”, “states” or “clusters” (referred to as “cell types” in the following) based on expression patterns and informed by prior knowledge. There are numerous methods to do this, including manually annotating clusters based on cell type markers, as well as automated “label-transfer” approaches that use annotated reference single-cell studies from the relevant tissue. Regenerating these cell type annotations from information provided in the publication can be difficult or impossible, due to the complexity and variability in the implementation of this step. For re-use, as well as for the purposes of reproducibility, it is of high value to have these annotations as determined by the authors of the study.

In 2021, (Puntambekar *et al*.) estimated that less than 25% of scRNA-seq studies deposited in GEO provide cell type annotations, based on data up to the end of 2020, and called for improved practices, but did not comment on processed data availability. Here we report on our recent experiences in expanding the scope of data types considered in Gemma to single-cell transcriptomic data, focusing on data released in GEO since 2021. We will not belabor considerations shared with all GEO studies, such as a lack of standardization of sample descriptions. Instead, here we focus on features of this particular data modality. Our findings confirm and extend the report of (Puntambekar et al., 2021) and reinforce the ongoing need for improving data sharing practices.

## Methods

To identify single-cell and single-nucleus datasets in GEO, using custom software (part of Gemma, https://github.com/PavlidisLab/Gemma), we first downloaded GEO’s metadata for 58,775 human, mouse, and rat transcriptomic experiments released between January 1, 2021 to December 31, 2024. A script was used to identify experiments containing the “Expression profiling by high-throughput sequencing” series type. The same script also screened for candidate single-cell transcriptomic experiments by screening for keywords in a case-insensitive manner in the metadata, including “scRNA”, “snRNA”, “single.cell”, “single.nuclei”, and “single.nucleus” (where “.” can be matched with any character). To reduce redundancy, SuperSeries were excluded if any of their SubSeries were present. The script accurately retrieved all 5,966 experiments annotated with ‘transcriptomic single cell’ as the Library Source, reflecting submitters’ designation of these datasets as single-cell studies.

Next, we downloaded supplementary data files available in GEO and analyzed them for the presence of processed data and cell type annotations. Filenames were used to determine whether files were associated with individual samples or the entire series, and to identify processed cell-level data provided in common formats such as the 10x Market Exchange (MEX). File formats were inferred from standard suffixes and extensions: *matrix.mtx. barcodes.tsv, features.tsv* or *genes.tsv* for MEX;.*rds*,.*RData*, or.*rda* for R-readable binaries; and.*h5ad* for AnnData. The contents of compressed files (.*zip*,.*gz*) and archives (.*tar*) were also inspected.

To look for candidate experiments that have cell-type annotations, we searched for cell-type containing files using keywords “coldata”, “meta”, “cell”, “annot”, “type” or “clustering” in the supplementary files. These files were then checked manually by curators.

Scripts and data supporting this paper are available at https://github.com/PavlidisLab/Persistent-Hindrances-to-Data-Re-use-in-Single-Cell-Genomics.

## Results

### Availability of processed data

We considered 58,775 human, mouse or rat experiments deposited in GEO between January 1st 2021 and December 31st 2024 (see Methods). By analyzing experiment metadata, we identified 13,037 candidate single-cell transcriptomic datasets. While the vast majority of these were transcriptomic experiments (library source is annotated as “transcriptomics” or “transcriptomic single cell”), 72 experiments of other types were identified and subsequently excluded (Figure 1a). Among the remaining studies, 5,966 were explicitly submitted to GEO as single-cell experiments (library source = “transcriptomic single cell”). However, the absence of this annotation does not necessarily indicate that a study is not single-cell.

**Figure 1:**
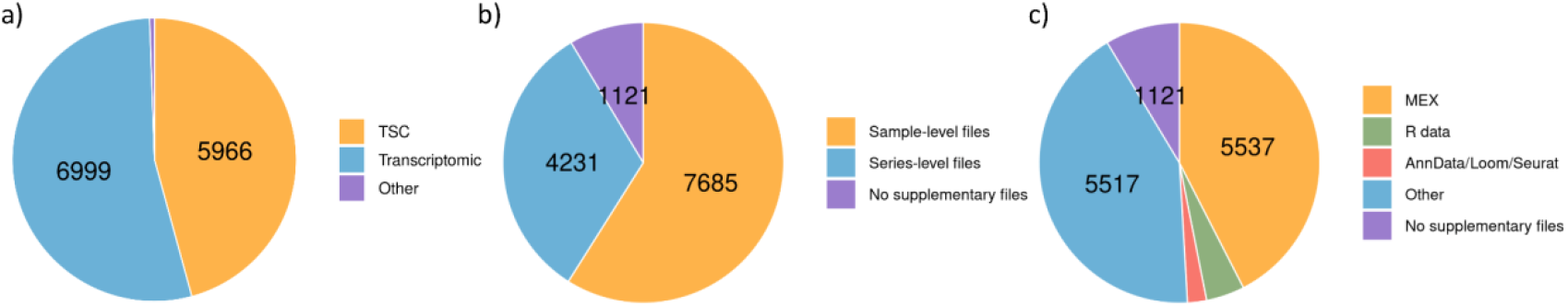
Characteristics of 13,037 candidate single-cell transcriptomic datasets.Experiment counts based on a) Library Source annotation, b) type of supplementary data, c) supplementary file formats.

To assess the availability of processed gene expression data, we first examined the names of the supplementary files provided. These filenames typically contain either a sample (GSM) or series (GSE) accession number, depending on whether the submitter associated the files with individual samples or with the overall series. It should be noted that files named with a GSE accession can still contain sample-level data—for example, when they are tar archives containing per-sample files, or when the filenames include sample descriptions. However, GSE-level files are difficult to systematically link to sample-level metadata within GEO records, whereas when files are provided per sample this is trivial. Across our corpus, the majority of datasets contained sample-level files (7,685/13,037; 59%), while 1,121 datasets lacked any supplementary files (Figure 1b).

We further examined supplementary files to identify commonly used formats for storing processed scRNAseq data (Figure 1c). The majority of experiments provided data in the 10x Market Exchange (MEX) format (5,537). In addition, 576 experiments included R-readable binary files, 246 were in AnnData format, 33 Loom and 7 Seurat. The remaining datasets either used less common formats or lacked supplementary files entirely (1,121).

### Availability of cell type annotations

We focused on studies with sample-level expression data in MEX format and retained only those with at least four samples (3,438), which makes them potentially suitable for differential gene expression analysis in Gemma. We used a custom script to search for specific keywords and narrow down experiments that might contain author-provided cell type labels in the GEO supplementary data files. This yielded 512 candidate experiments, which we then manually reviewed. Of these, we determined that only 265 actually contained cell-type annotations, representing just 8% (265/3438) of the total studies considered (Figure 2).

**Figure 2:**
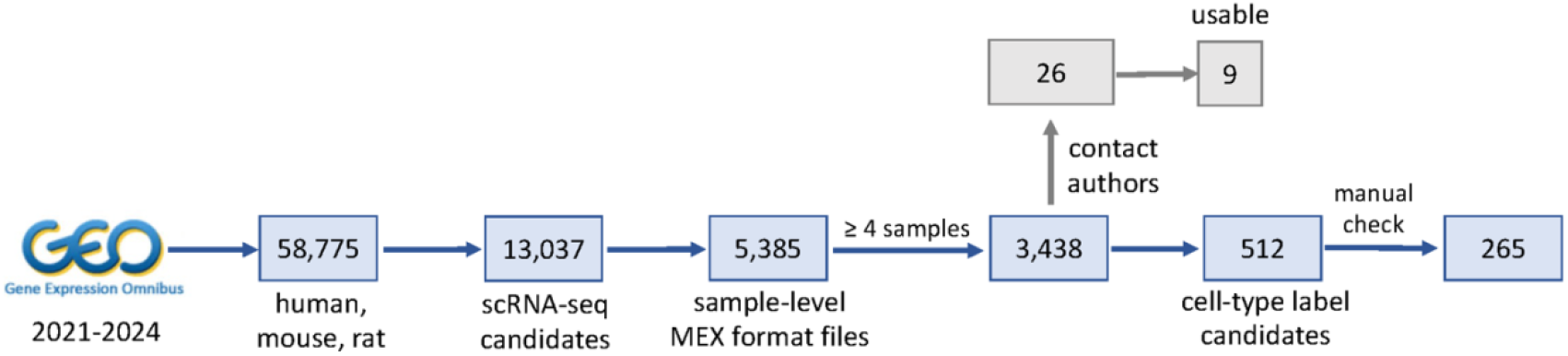
Flowchart of the GEO experiment screening process.

Next, we tested whether missing cell type annotation files could be found in the associated publications or obtained from the authors, for a sample of 27 brain-related studies (prioritized for Gemma) which lacked annotations in the GEO record. Only one had cell-type annotations provided as a supplementary data file in the associated publication. We then wrote to the corresponding authors of the remaining 26 studies requesting the data, attempting to reach each three times. Six explicitly denied our request on various grounds (discussed below). Eight failed to respond. The remainder (12, 46%) provided the requested files. However, three of the provided data files were not usable (discussed below), leaving a final yield of 9, or 35% of those contacted.

The seven denials of our requests had various justifications. In three cases, the authors told us to do the cell type annotations ourselves, since the methods were described in the paper. In five cases, the data was reported as lost, and in one case the authors said they would have to regenerate the annotations but were too busy. Author-provided cell annotation files which were not usable had a variety of problems. The most common issue was the cell identifiers could not be matched to those in the data provided in GEO, rendering the data useless. We also encountered a case where many cells were annotated to multiple cell types, suggesting some error that we could not resolve.

### Lack of consistency in cell annotations

Focusing on the studies that had cell type information provided (265+12), we further explored the quality of these annotations. Cell types were never (0%) labelled with formal identifiers such as ontology terms, instead all being free text. Even for commonly recognized cell types such as microglia (resident macrophages of the central nervous system), authors referred to them with a variety of terms such as “MGs”, “Micro”, “MicroG”, etc. (note we are inferring from the context that these abbreviations are for “microglia”; to be clear, no key to the terms is ever provided by the submitters). Some datasets provide uninformative cell type labels such as “Cluster 1” (e.g. in GSE132355). For such datasets and in other cases, mapping to ontology terms or other consistent terminologies would be difficult or impossible.

Often, sample identifiers provided in the supplementary files do not match those in the GEO metadata or require a certain degree of transformation, making it difficult to assign individual cells and their annotations to the correct sample. In some cases, we were able to use cell barcodes to make the link, but this fails if there are any collisions (barcodes assigned to more than one cell).

### Trends over time

We hypothesized that over the period sampled, there might be some improvement in practices. We therefore used the GEO study release dates to explore trends in data submission quality over time. At least one improvement is evident. Among the candidate single-cell experiments (13,037), we found that the proportion tagged with “transcriptomic single cell” as the Library Source has steadily increased, from 1% to 70%, (Figure 3), making it easier to reliably identify single-cell experiments in GEO. Similarly, there is an upward trend for the deposition of the more standardized cell-level expression file format: the proportion of experiments with files in MEX format climbed from 31% to 46% during the past four years (Figure 4). We see the opposite trend when it comes to cell-type annotations: among the 3,438 candidate experiments, the proportion with submitter-provided labels dropped from 8.9% to 6.4% over the past four years (Figure 5).

**Figure 3:**
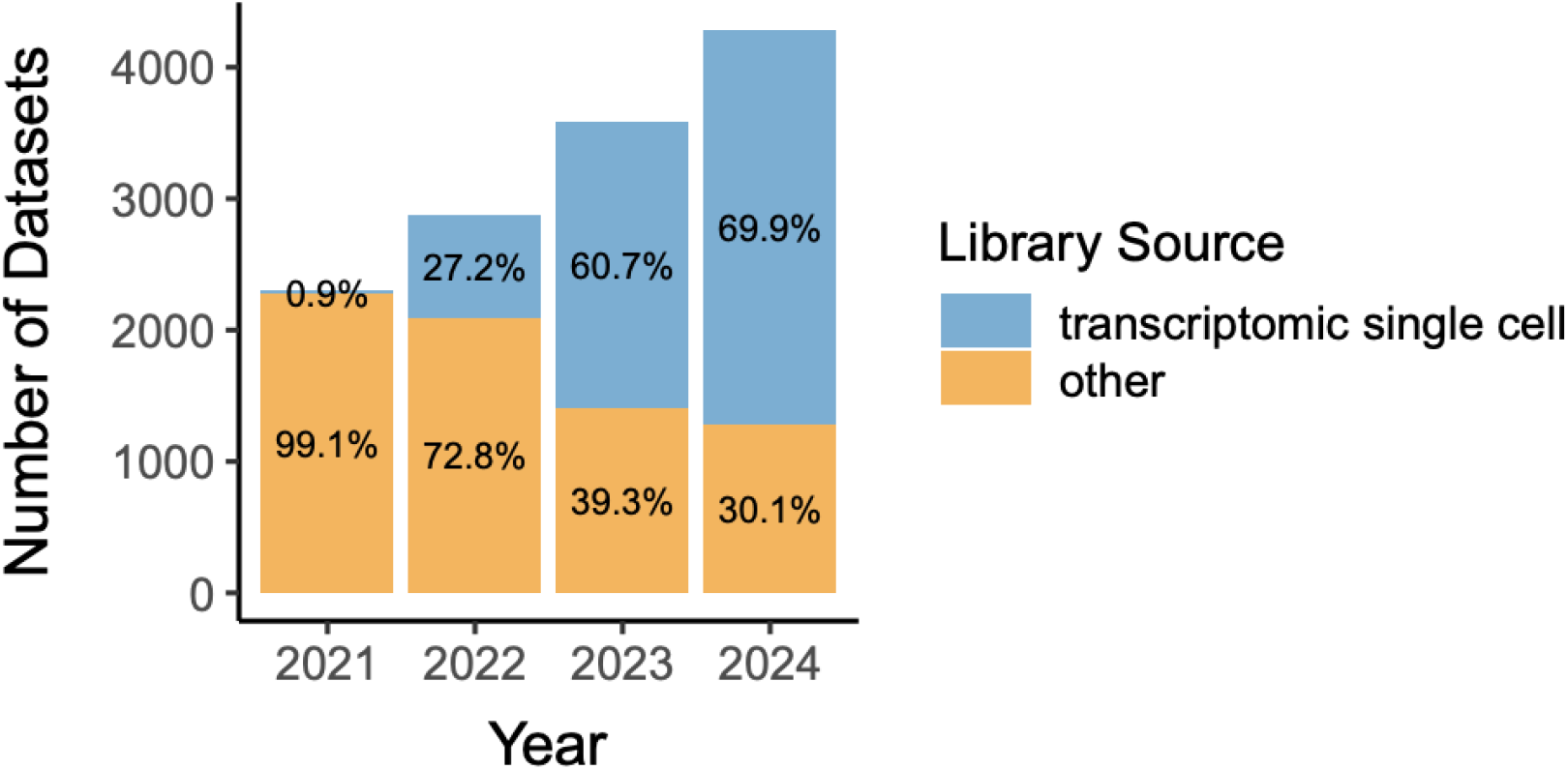
The use of the formal Library Source annotation “Transcriptomic single cell” has increased over time. Data are shown for the 13,037 candidate single-cell GEO studies submitted between 2021 and 2024. The datasets were selected by searching GEO metadata for “Expression profiling by high-throughput sequencing” and keywords indicative of single-cell focus of the study.

**Figure 4:**
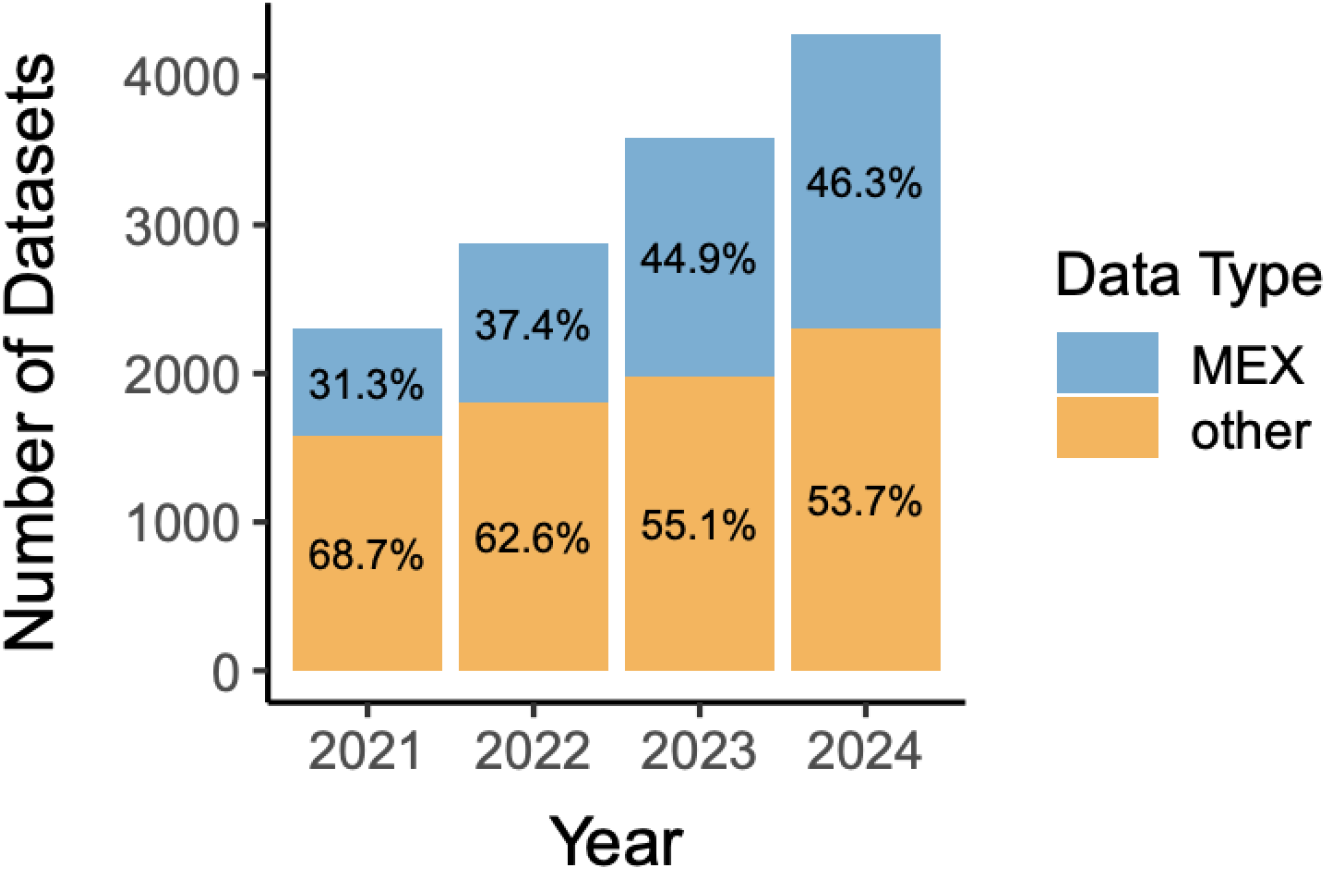
The provision of processed cell-level gene expression data files in the standard 10x Market Exchange (MEX) has increased over time. Data are shown for the 13,037 candidate single-cell GEO studies submitted between 2021 and 2024. The files were identified by examining file names, extensions and the contents of archives (TAR and ZIP) among the supplementary materials available in GEO. Only files containing data at the sample level were considered.

**Figure 5:**
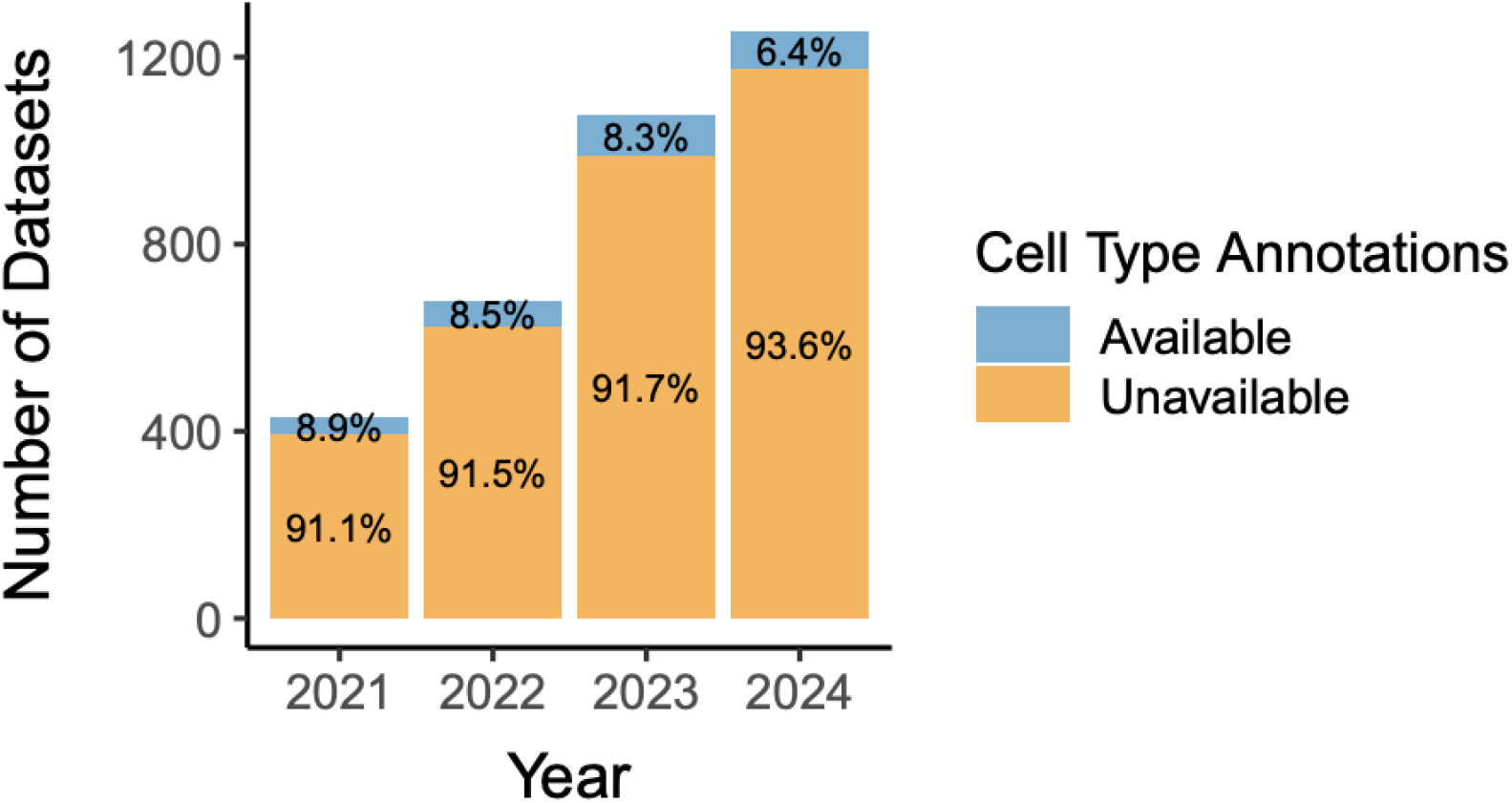
The provision of cell-type annotations has decreased over time. Data are shown for the 3,438 candidate single-cell GEO studies submitted between 2021 and 2024. The candidate studies were selected to include four or more samples and have processed cell-level expression data in MEX format. The studies were first programmatically screened for keywords of interest and then manually inspected to confirm the presence of cell-type annotations.

## Discussion

Our survey of single-cell transcriptome studies in GEO shows that current community standards for data submission are below what would be required for routine re-use of the data. These results suggest that the concerns raised by (Puntambekar *et al*., 2021) based on data available at the end of 2020 have not improved, and may have in fact gotten worse in some ways.

The lack of readily usable processed gene-expression data for more than half of GEO single-cell submissions is an issue shared to some extent with other RNA-seq modalities in GEO, but has not been previously reported. Firstly, within our corpus of 13,037 datasets, 1,121 lacked supplementary files altogether. Processed scRNA-seq data files are generally not provided as supplements to publications, meaning one would have to reach out to the authors or re-analyze the raw FASTQ files (if available) to obtain expression levels. Even when author-supplied supplementary files are present, they are not always in an easily usable format. For bulk RNA-seq, processed data are conventionally represented as a single gene-by-sample count matrix. By contrast, single-cell studies introduce an additional dimension (gene × sample × cell), increasing the complexity of data representation and resulting in greater heterogeneity in file formats. While some formats, such as MEX and AnnData, have been adopted as standards, GEO submissions continue to include processed data in a wide variety of non-standardized formats. Furthermore, series-level data files (filenames containing GSE accessions), even when they contain sample-level count data, are often difficult to associate with the corresponding sample-level metadata. Consequently, our automated Gemma workflow could only incorporate experiments with sample-level count data in MEX format, which is only 41% (5385/13037) of the initial corpus(Figure 2).

GEO does not provide expression level information for RNA-seq datasets in their core data model, though they have started recomputing them from FASTQ files for some studies, but this has not yet been extended to single-cell data. Relying on author-submitted supplementary files, which are not fully validated by GEO, leaves the potential for non-adherence to guidelines and other issues. In contrast, for microarray studies, GEO imports expression levels as structured data. Sometimes, microarray supplementary files are easy to use, as the case for Affymetrix GeneChip data where CEL files are usually provided in GEO and can be easily re-analyzed, as we do for Gemma. On the other hand, the Sequence Read Archive (SRA) provides raw data for re-analysis of GEO RNA-seq datasets from FASTQ files. Reanalyzing these data is possible and can be automated (as in Gemma). For single-cell studies on the 10X platform, the need to use the proprietary Cell Ranger software is a relatively small hurdle. However, without complete details of how the software was applied, reproducing specific findings can be hindered by differences in software version, reference transcriptome, or analysis parameters. Moreover, we expect the re-processing of SRA scRNA-seq data to be more challenging than for “standard” RNA-seq data, where FASTQ files are provided de-multiplexed and the relationship of files to GEO samples is generally straightforward.

While we have not yet fully evaluated the ease of use of SRA files to fill the gap for scRNA-seq, we carried out preliminary analyses focused on raw data availability and usability. Within our corpus of 13,037 datasets, 960 lacked any SRA record and of the remaining 11,708 transcriptomic datasets, 731 had no valid runs. Only 2,036 datasets were associated with an SRA record containing complete metadata, while the remaining 8,941 datasets had metadata deficiencies that can complicate or prevent reprocessing. A common problem was the absence of a loader script (fastq-loader.py) linking author-submitted FASTQ files to the SRA metadata (5,708 datasets). In such cases, we must resort to heuristics to infer associations, such as matching file sizes with read lengths.

In our view, the lack of cell-type annotations is even more problematic. Our estimate for the number of datasets with cell type annotations is substantially below Puntambeker et al.’s estimate of “<25%” - we believe it is less than 10% (our point estimate is 8%). The cell type assignments are usually a crucial component of any publication using single-cell data, forming the basis of the interpretation and statistical analysis of the data. We foresee that failing to address this will mean sustained hurdles to replicating findings and re-using the data. Contacting authors for cell type annotations proved to be moderately effective, if time-consuming. If our yield of 35% is representative, we suggest that this path is worth pursuing, but not a solution. Alternatively, computational reannotation of cell types is becoming easier, though this requires accepted “references”, which are not yet available for every tissue and developmental stage. Furthermore, aligning re-computed annotations with the particularities of studies where the authors (for example) manually split a “recognized” cell type into multiple *ad hoc* subclasses (e.g., oligodendrocytes 1, oligodendrocytes 2) means replicating the original results will be nearly impossible.

Since 2021, when Puntambekar et al. reported on some of the same issues, GEO has revised its submission requirements to include a dedicated section for single-cell studies and has adopted more prominent and specific language regarding the deposition of processed scRNA-seq data. However, despite the stated requirement for processed data, enforcement remains inconsistent - many datasets are still submitted without it. In addition, while the annotation of single-cell experiments with “transcriptomic single cell” as the Library Source has improved, it remains inconsistent. This label is automatically assigned by GEO when submitters use a specific Library Strategy tag for single-cell RNA-sequencing samples (“scRNA-seq”) in the GEO metadata submission spreadsheet. However, we find that up to 30% of single-cell datasets still lack this annotation (Figure 3).

Additionally, the updated GEO guidelines still do not require the inclusion of cell-type annotations. Authors should be advised to provide clear descriptions of the cell types they annotated, ideally using ontology terms or at least something better than “Cluster 1”. We note that CellXGene (CZI Single-Cell Biology Program et al., 2023) has already implemented these requirements, but they have a narrower focus and less data than GEO at this time. It may be too late to do anything about the data that is already in GEO, but firming up requirements now will pave the way for more efficient re-use of data in the future.

We acknowledge the utility of flexibility for scientists to cluster cells in a way that meets their needs, so “microglia 1” and “microglia 2” may be appropriate in a particular study. It is especially in such cases of “bespoke” cell type annotation that providing the data is critical, especially if the annotation was done manually or iteratively in a way that can’t be computationally reproduced. But when it comes to re-use, such flexibility can seem like chaos. We suggest a hybrid approach, where ontology terms are used as much as possible and study-specific features like “cluster 2” can be added on top of that.

In summary, the current GEO submission practices for single-cell RNA-sequencing data remain inadequate for effective and routine re-use. Strengthening submission requirements — particularly around processed data and cell-type annotations — and enforcing them consistently will be essential to realize the full value of these increasingly prevalent datasets. But for the moment, re-users of data will often have to start over from scratch from FASTQ files (if available), or rely on resources such as Gemma that apply consistent processing and annotation pipelines.

## Acknowledgements

We thank the thousands of researchers who have submitted data to GEO, making our work possible.

## Author contributions

PP: Designed the study, directed work, contributed to manuscript development. SR: Directed work, drafted the manuscript. RS: Contributed to study design and data interpretation. XXY, BX, AM, SS, G-PM: Data generation, analysis and curation.

## Funding

This work was supported by National Institutes of Health grant MH111099 and Natural Sciences and Engineering Research Council of Canada (NSERC) grant RGPIN-2016-05991 held by PP.

## Notes

### Competing Interest Statement

The authors have declared no competing interest.

### Summary of Updates

Fixing two minor errors in the text

